# MAP1LC3C regulates lysosomal exocytosis and induces zinc reprogramming in renal cancer cells

**DOI:** 10.1101/2022.05.11.491473

**Authors:** Rita Verma, Parul Aggarwal, James Reigle, Dina Secic, Collin Wetzel, Megan E. Bischoff, Katherine VandenHeuvel, Jacek Biesiada, Birgit Ehmer, Julio A. Landero Figueroa, David R. Plas, Mario Medvedovic, Jarek Meller, Maria F. Czyzyk-Krzeska

**Affiliations:** Department of Cancer Biology, University of Cincinnati College of Medicine, Cincinnati, OH 45267, USA; Department of Biomedical Informatics, University of Cincinnati College of Medicine, Cincinnati, OH 45267, USA; Division of Biomedical Informatics, Cincinnati Children’s Hospital Medical Center, Cincinnati, OH 45229, USA; Division of Pathology and Laboratory Medicine, Cincinnati Children Hospital Medical Center, Cincinnati, OH, USA; Division of Biostatistics and Bioinformatics, Department of Environmental and Public Health Sciences University of Cincinnati College of Medicine, Cincinnati, OH 45267, USA; Agilent Metallomics Center of the Americas, Department of Chemistry, University of Cincinnati College of Arts and Science, Cincinnati, OH 45221, USA; Department of Electrical Engineering and Computer Science, University of Cincinnati College of Engineering and Applied Sciences, Cincinnati, OH 45221, USA; Veteran Affairs Medical Center, Department of Veterans Affairs, Cincinnati, OH 45220, USA; Department of Pharmacology and System Biology, University of Cincinnati College of Medicine, Cincinnati, OH 45267, USA

**Author notes:** Contact author: Maria F. Czyzyk-Krzeska, MD, PhD., Department of Cancer Biology, University of Cincinnati, College of Medicine, 3125 Eden Ave, Cincinnati, OH 45267. Trace Elements Group, Department of Environmental Medicine and Public Health at Icahn school of Medicine at Mount Sinai, 1428 Madison Ave, NYC, NY-10029.

## Abstract

MAP1LC3C (LC3C) is a member of the microtubule associated family of proteins that are essential in the formation of autophagosomes and lysosomal degradation of cargo. LC3C has tumor suppressing activity and its expression is dependent on kidney cancer tumor suppressors, such as VHL and FLCN. Recently we demonstrated that LC3C autophagy is regulated by noncannonical upstream regulatory complexes and targets for degradation postdivision midbody rings associated with cancer cells stemness. Here we show that loss of LC3C leads to peripheral positioning of the lysosomes and lysosomal exocytosis (LE) in a subset of cells. This process is independent of the autophagic activity of LC3C. Analysis of isogenic cells with low and high LE shows substantial transcriptomic reprogramming with altered expression of Zn-related genes and activity of Polycomb Repressor Complex 2 (PRC2), accompanied by a robust decrease in intracellular Zn. Metabolomic analysis revealed alterations in amino acid steady-state levels. Cells with augmented LE show tumor initiation properties and form aggressive tumors in xenograft models. Immunocytochemistry identified high levels of LAMP1 on the plasma membrane of cancer cells in human ccRCC and reduced levels of Zn, an indication that LE is a frequent event in ccRCC, potentially contributing to the loss of Zn. Overall, these data indicate that an important tumor suppressing activity of LC3C is contributing to the reprogramming of lysosomal activity and Zn metabolism with implication for epigenetic remodeling in a subpopulation of tumor propagating properties of cancer cells.

## Introduction

Lysosomes function as the final organelles destined to degrade cargo delivered by autophagosomes and multivesicular bodies. They occupy primarily a perinuclear region of the cell in proximity to microtubule organizing centers. However, lysosomes can also move away from the nucleus in response to stimuli and an important activity is the secretion of their content into the extracellular environment, a process called lysosomal exocytosis (LE) (1). During LE, lysosomes translocate from their perinuclear localization into to the proximity of cell surface via the microtubule-dependent motor protein kinesin. Then they fuse with the plasma membrane in a Ca^2+^ dependent manner, release their luminal content to the exterior of the cell and expose the luminal site of their integral lysosomal proteins, such as LAMP1, on the extracellular aspect of the plasma membrane (2). LE is transcriptionally regulated by TFEB transcription factor, a master regulator of genes in lysosomal function (3). TFEB stimulates expression of genes involved in the tethering and docking of the lysosome to the plasma membrane. Moreover, TFEB induces expression of lysosomal transient receptor potential mucopolin 1 channel (TRPML1/MCOLN1), which releases lysosomal calcium triggering the final fusion (4). LE allows for cellular clearance and detoxification, including toxic levels of metals (4-6), excretion of active proteases that remodel extracellular matrix and contribute to cell motility (7, 8). LE is necessary for membrane repair and remodeling that occur during phagocytosis and neurite outgrowth (9, 10). While release of exosomes is usually attributed to the multivesicular bodies and their fusion with the plasma membrane, LE can also release certain types of exosomes (1). Thus, LE affects paracrine intercellular communications in multiple manners.

LE plays important role in cancer, promoting cancer progression due to the remodeling of extracellular matrix and modulation of tumor cell invasion, angiogenesis and metastasis. LE was shown to play a role in progression of sarcomas and gliomas (8, 11). In particular, high levels of LAMP1, a sialylated glycoprotein in the lysosomal membrane promote LE, a process that can be reversed by the activity of NEU1, sialidase that prevents oversialylation of LAMP1 (12). LE plays also important role in chemoresistance (12, 13). Lysosomal sequestration of chemotherapeutics induces LE preventing them from reaching their intracellular targets (13).

MAP1LC3C (LC3C) is a paralog in the microtubule-associated protein light chains family of LC3 autophagic regulators. LC3s become lipidated on the C-terminal glycine, which allows for the insertion into the elongating membrane of the forming autophagosome and tethers the cargo receptors bringing ubiquitinated cargo to the autophagosomes. LC3C autophagy is an evolutionarily late program present only in higher primates, including humans. LC3C differs in amino acid sequence with other paralogs by 60% and contains a 20 amino acid long C-terminus peptide located after the Glycine 126 which undergoes lipidation, absent in other paralogs. Compared to LC3C, the LC3A and LC3B paralogs form separate autophagic vacuoles supporting different functions (14). LC3C is induced by the von Hippel Lindau tumor suppressor (VHL), in the mechanism that involves repression of the hypoxia inducible transcription factor (15). LC3C acts as a tumor suppressor in ccRCC, where it represses accumulation of postdivision midbodies, markers of cancer cell stemness, indicating that the tumor suppressing activity of LC3C limits the activity of cancer stem cells (14). LC3C was also shown to have tumor suppressing activity in breast cancer, where it contributes to degradation of Met receptor tyrosine kinase and regulates its downstream signaling and cell migration and invasion (16). Recently, we reported that LC3C autophagy is regulated by the non-canonical preinitiation and initiation complexes that include ULK3, UVRAG, Rubicon and PIK3C2A (14).

Here we show that loss of LC3C results in the translocation of lysosomes towards cell periphery and activation of lysosomal exocytosis in a subset of cells. This process does not involve autophagic activity of LC3C. Transcriptomics comparison of isogenic cells with low and high LE demonstrate robust reprograming related to the activity of Polycomb Repressor Complex 2 (PRC2), major decrease in the intracellular Zn and enrichment for the altered expression of Zn-related genes. Metabolomic analysis revealed also major changes in amino acid steady-state levels. Cells with augmented LE show tumor initiation properties and form aggressive tumors in xenograft models. Immunocytochemistry identified high levels of LAMP1 in the plasma membrane of cancer cells in human ccRCC and reduced levels of Zn, an indication that LE is a frequent event in ccRCC, potentially contributing to the loss of Zn. Overall, these data indicate that an important tumor suppressing but not autophagic activity of LC3C is regulation of lysosomal activity and Zn metabolism, suggesting epigenetic remodeling in a subpopulation of tumor propagating properties of cancer cells.

## Results

### Loss of LC3C induces LE in autophagy-independent manner

Positioning and motility of lysosomes are regulated by cancer driver events, environmental factors and nutritional conditions, including amino acid availability, hypoxia or cholesterol load (17). Investigating the effects of VHL in clear cell renal cell carcinoma (ccRCC) cells, we previously established that reconstitution of VHL leads to de-repression of LC3C transcription through degradation of HIFs (15). Knock-down of LC3C leads to formation of tumors in orthotopic xenografts, establishing the tumor suppressing activity of LC3C in ccRCC (15). Interestingly, stable knockdown of LC3C in 786-O VHL(+) induced significant repositioning of lysosomes characterized by dispersed LAMP1 and CTSD staining from the perinuclear region, throughout the cell towards plasma membrane (Fig. 1A and 1B), suggesting increased lysosomal exocytosis. Consistent with this hypothesis, we determined the presence of the lysosomal luminal domain of LAMP1 on the plasma membrane surface in cells with LC3C-KD, when immunofluorescence analysis was performed on cells that were not permeabilized (Fig. 1C). Further on, flow cytometry measured a significant increase in the number of cells with plasma membrane localization of LAMP1 in 786-O VHL(+) and Caki-1 cells with LC3C KD (Fig. 1D and 1E). Pretreatment of LAMP1^*H*^ cells with lysosomal inhibitors, vacuolin and chloroquine, for 24 h significantly diminished number of cells with LAMP1 on plasma membrane (Fig. 1F). Next, we determined that media from cells with LC3C, but not LC3B, knocked down showed accumulation of cleaved cathepsin D (CTSD) heavy chain, however, levels of procathepsin D were less affected (Fig. 1G, top). This is an indication of the release of the active lysosomal enzymes rather than cathepsin D precursors secreted in exosomes. There was also an increase in the cathepsin levels in the cellular lysates (Fig. 1G, bottom). Finally, we found that increased presence of LAMP1 on cell surface and release of active cathepsin into the extracellular media measured in cells with LC3C-KD were reversed (Fig. 1H and 1I) by reexpression of wild type LC3C, and also the autophagy deficient LC3C mutant, G126A (Fig.1J). This indicates an important, autophagy independent function of LC3C in trafficking of lysosomes to the plasma membrane.

**Figure 1.**
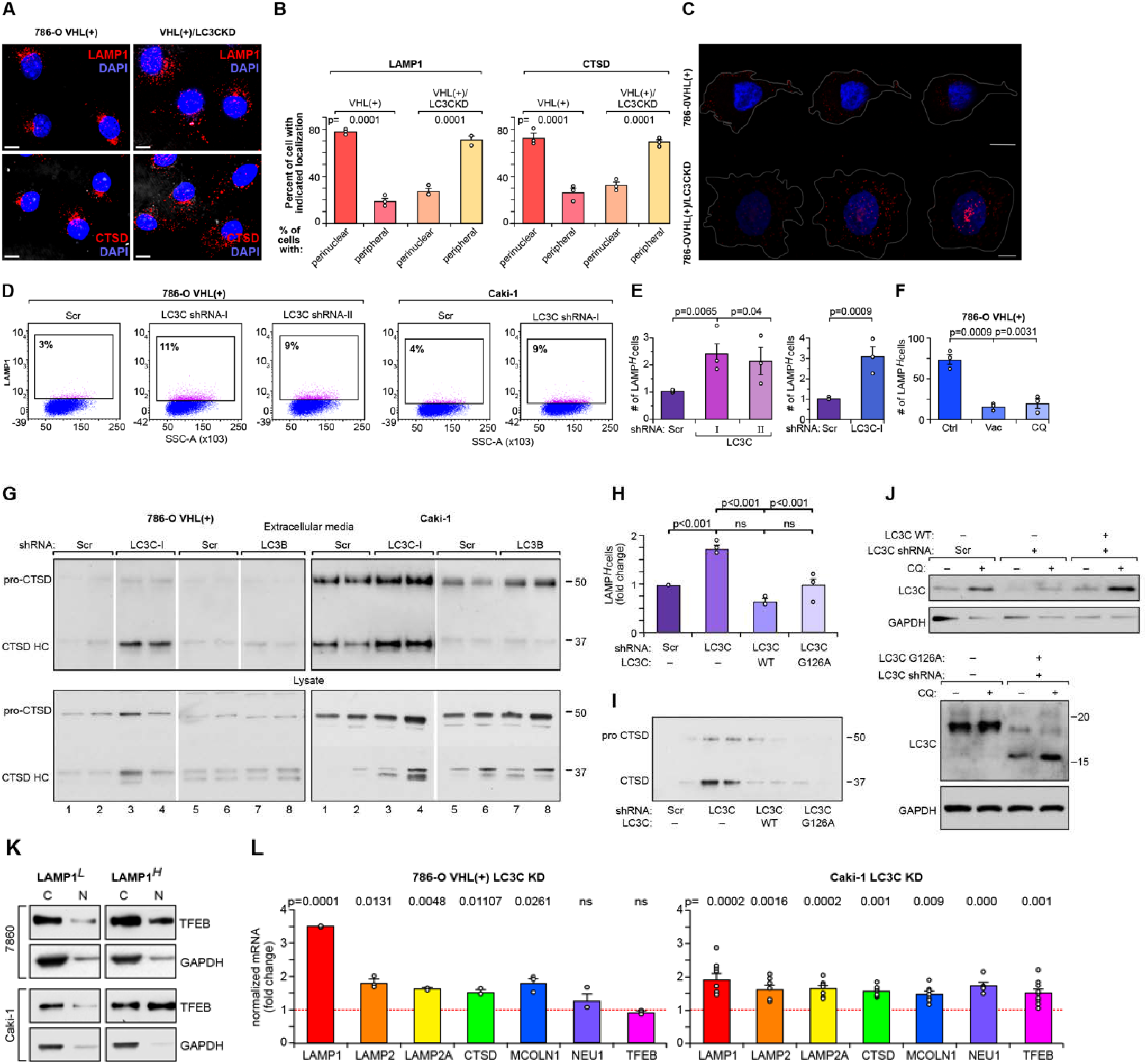
Loss of LC3C induces LE: (A) Immunofluorescence for LAMP1 and CTSD in 786-0 VHL(+) cells with LC3C-KD. Scale bar = 10 µM; (B) Quantification of perinuclear vs peripheral distribution of lysosomes using LAMP1 and CTSD markers. (C) Z-stack showing Immunofluorescence of LAMP1 on the surface of non-permeabilized cells with indicated status of LC3C. Outlines of the cells are shown. Scale bar =10 µM; (D) Scatter plots showing increased number of cells with surface localization of LAMP1 in response to LC3C-KDs in the indicated cell lines. (E) Quantification of enrichment for cells with LAMP1 localized to plasma membrane in the indicated cell lines. (F) Quantification of cells with LAMP1 localized to plasma in cells treated with lysosomal inhibitors vacuolin-1 (Vac, 5µM) and chloroquine (CQ, 20 µM) for 24 h. (G) Western blots show pro-CTSD and cleaved CTSD heavy chain (HC) in the concentrated tissue culture media (top) and cellular lysates (bottom) in the indicated cell lines. (H) Effects of reexpression of LC3C wild type (WT) or autophagy deficient mutant (G126A) in cells with LC3C-KD on the number of cells with plasma membrane localization of LAMP1. (I) Effects of reexpression of the LC3C constructs of accumulation of pro-CTSD and CTSD HC in extracellular media. (J) Western blots show expression of exogenous LC3C construct similar to the expression of endogenous LC3C. (K) Western blots show translocation of TFEB from the cytosolic to the nuclear fraction. (L) Results of qRT-PCR show induction of mRNA expression for several lysosomal proteins in the indicated cell lines. All p values obtained by two-tailed t-test.

Next, we sorted 786-O VHL(+) and Caki-1 cells with stable LC3C-KD cells in order to enrich for cells expressing or not expressing LAMP1 on the plasma membrane (LAMP1^*H*^ vs. LAMP1^*L*^). LE is induced at the transcriptional level by the master transcription factor, TFEB, that induces expression of signature of lysosomal and autophagic genes (3). Consistently, we found nuclear translocation of TFEB (Fig. 1K) and significantly increased gene expression for LAMP1, LAMP2, CTSD and MCOLN1 in LAMP1^*H*^ as compared to LAMP^*L*^ cells selected by flow cytometry (Fig. 1L). NEU1 and TFEB mRNA expression was increased only in Caki-1 but not 786-O cells (Fig. 1L). These data indicate induction of lysosomal biogenesis in cells with LC3C KDs with enriched LE.

### High LE activity corresponds with transcriptional responses to reduced Zn content

In order to determine the molecular effects triggered by LE, we performed RNA-seq analysis of RNA extracted from four biological replicates of LAMP1^*H*^ and LAMP1^*L*^ cells isolated by FACS. Unsupervised clustering using differentially expressed genes and Pearson correlation-based distance measure clearly stratified the samples by LAMP1 expression (Fig. 2A). We found 1096 genes expressed at two-fold difference and FDR<0.01 (Supplemental Table 1). 566 genes were upregulated and 530 gene were downregulated. There was a significant enrichment for the Zn-related genes among genes differentially regulated in LAMP1^*H*^ and LAMP1^*L*^ cells as compared to the proportion of these genes in RefSeq database (Fig. 2B, Supplemental Table 2). Direct measurement of intracellular Zn using size exclusion chromatography coupled with inductively coupled plasma mass spectrometry (SEC-ICP-MS) revealed decreased total Zn content as well as in protein bound-Zn, corresponding to high molecular weight fraction (HMW), metallothioneins (MT), small cysteine rich proteins that buffer metals, and Zn present in the low molecular weight fraction (LMW) in LAMP1^*H*^ cells (Fig. 2C). This is likely related to the function of lysosome serving as Zn storage and the role of LE as a mechanism for Zn excretion (6, 18). This indicates that LE by altering cellular Zn pools leads to global reprogramming, contributing to its effects in tumor progression.

**Figure 2.**
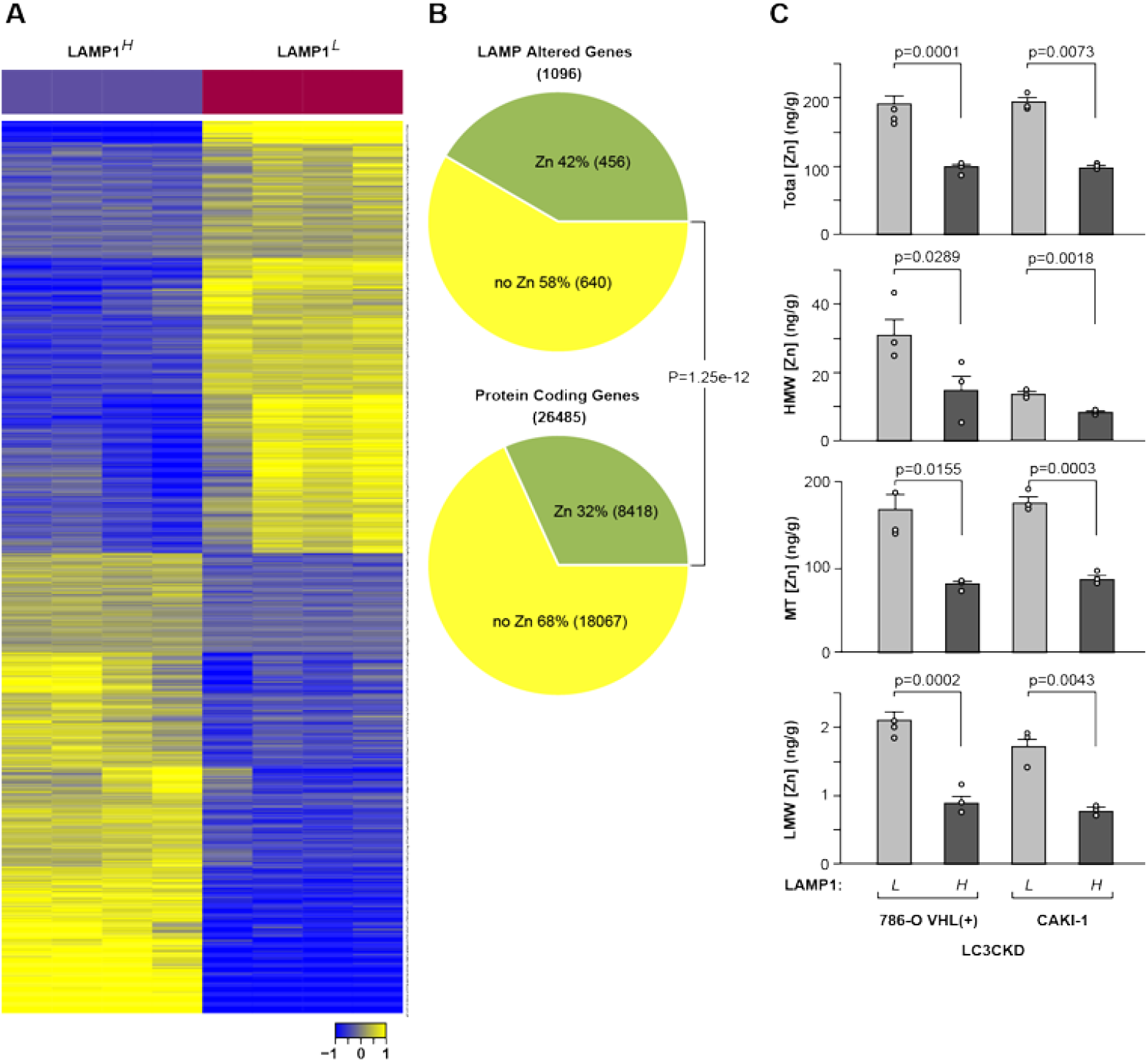
Cells with high LE show altered expression of Zn-related genes and decreased levels of Zn in several molecular fractions. (A) Heatmap shows differential gene expression between LAMP1^*H*^ and LAMPL^*L*^ cells. (B) Pie chart shows significant enrichment for Zn-related genes among genes differentially expressed between LAMP1^*H*^ and LAMPL^*L*^ cells as compared to the number of Zn-related genes among all protein coding genes in RefSeq database. P value from chi-square proportions test. (C) Measurement of the Zn concentration in total cellular lysates (top) and in the high molecular weight fraction (HMW), Zn bound to metallothioneins (MTs) and Zn present in low molecular weight fraction (LMW) in LAMP1^*H*^ and LAMPL^*L*^ cells enriched by sorting in the indicated cell lines. P values obtained by two-tailed t-test.

### LE leads to Zn-dependent epigenetic reprogramming

Analysis of transcription factors related to the genes differentially regulated between in LAMP1^*H*^ and LAMP1^*L*^ cells in Enrichr revealed a significant enrichment of binding sites specific for the SUZ12 member of Polycomb Repression Complex 2 (PRC2) using ENCODE and CHEA database (Fig. 3A), and for histone 3 K27 trimethylation by Epigenomics Roadmap HM ChiP-seq database (Fig. 3B). This is consistent with PRC2 activity as H3K27 methyltransferase maintains epigenetic repression of silenced genes. Importantly, 95 genes upregulated in LAMP^*H*^ cells were associated with SUZ12 and EZH2 in ENCODE and ChEA Consensus TF and ENCODE TFChIP-seq 2015, and with H3KMe3 in ENCODE Histone Modifications 2015 and Epigenomics Roadmap HM ChIP-seq databases (Fig. 3C, Supplemental Table 3). The most significant pathway enriched among these genes was Epithelial Mesenchymal Transition (MSigDB Hallmark 2020).

**Figure 3.**
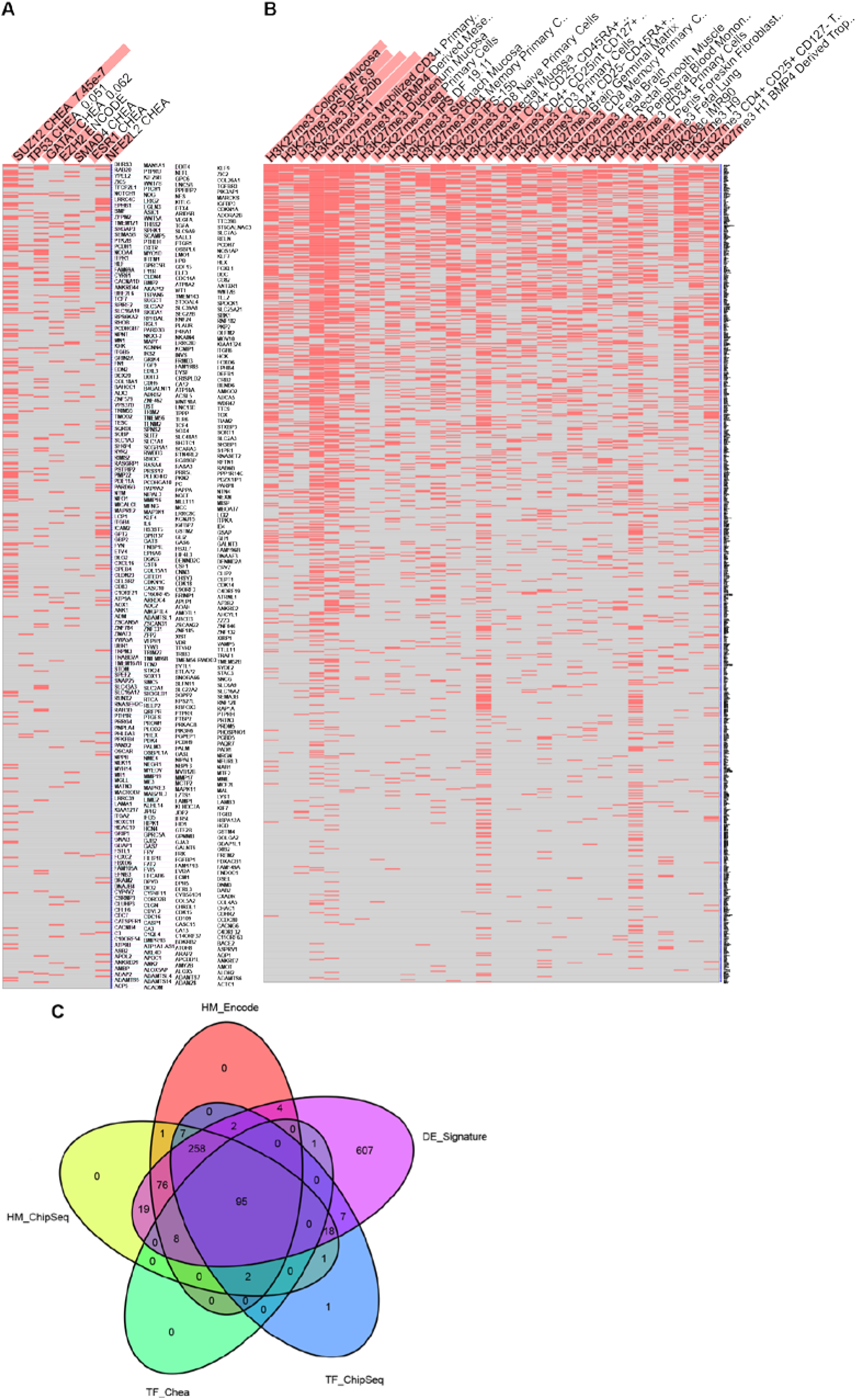
Analysis of transcriptional regulators common for the altered Zn-related genes shows enrichment for components of PRC2 complex. (A) Enrichr analysis using ENCODE and CHEA database shows enrichment for SUZ12 and EZH2 transcription factors for genes differentially expressed in LAMP1^*H*^ and LAMPL^*L*^ cells. (B) Enrichr analysis using Epigenomics Roadmap HM ChiP-seq database shows enrichment for H3K27Me3 for genes differentially expressed in LAMP1^*H*^ and LAMPL^*L*^ cells. (C) Venn diagram identifies 95 genes differentially expressed in LAMP1^*H*^ and LAMPL^*L*^ cells and determined to be regulated by SUZ12/EZH2 and H3K27Me3 in the indicated databases.

Fractionation of cell lysates into nuclear and chromatin fractions revealed overall decrease in LAMP1^*H*^ cells of H3K27Me3 and components of PRC2, SUZ12, EZH2 and MTF2 while there was an increase in H3K27Ac (Fig. 4A-4D). This indicates major PRC2-related transcriptional reprogramming. PRC2 subunits utilize Zn ions for their activity. SUZ12 has essential for its activity C2H2-type Zn finger motif (19). EZH2 has CXC domain with two Zn3Cys9 motifs in its SET domain (20). Moreover, the polycomb-like (PCL) protein necessary for recruitment of PRC2 to the nucleation sites, PCL2 or MTF2, is also a Zn-finger transcription factor binding to metal responsive elements regulating expression of the MTF2 gene (21). Consistently, expression of MTF2 mRNA was decreased in LAMP1^*H*^ cells (Fig. 4E). We also determined that there was a significant decrease in the Zn concentration in the chromatin fraction (Fig. 4F), supporting Zn-responsive epigenetic reprogramming.

**Figure 4.**
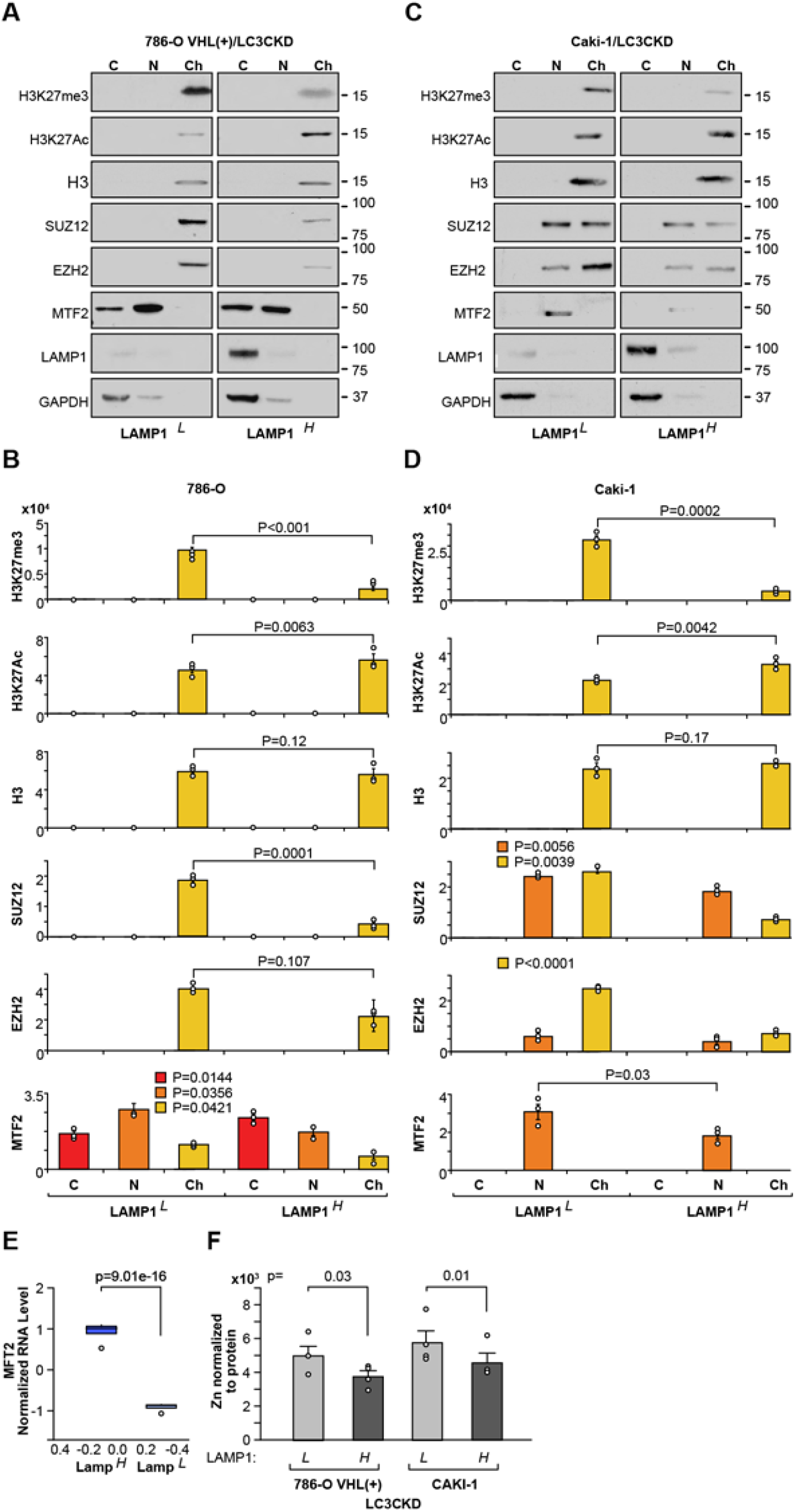
Cells with increased LE show diminished H3K27me3 and decreased presence of PRC2 components in chromatin fraction. (A & B) Western blots and (C & D) respective quantification for H3K27me3, H3K27 Ac and indicated components of PRC2 complex. (E) analysis of expression of MTF2 mRNA expression in LAMP1^*L*^ and LAMP1^*H*^ cells. (F) measurement of Zn concentration in the chromatin enriched fraction as shown in A and B. P values obtained by two-tailed t-test.

### Activation of LE leads to metabolomic reprogramming

Metabolomic analysis of cellular lysates revealed clear differences in the steady-state levels of 77 metabolites at FDR<0.05 between LAMP^*H*^ and LAMP^*L*^ cells (Fig. 5A, Supplemental Table 4). Levels of 31 metabolites were higher in LAMP^*H*^ cells, while levels of 46 metabolites were higher in LAMP^*L*^ cells. There was a highly significant enrichment for pathways related to amino acid metabolism and protein synthesis (Fig. 5B and 5C). In particular, steady-state levels of most amino acids, including glutamate, valine, serine, threonine, tyrosine, histidine, alanine, tryptophan, leucine, glycine, aspartate, proline and glutamine and methionine were decreased in LAMP^*H*^ cells (Fig. 5D). Only levels of three amino acids, arginine, lysine and cysteine, were increased in LAMP1^*H*^ cells (Fig. 5E). This overall decrease in amino acid levels in LAMP1^*H*^ cells can be attributed to the diminished activity of lysosomes in cellular biodegradation of protein and regeneration of nutrients because of their destination towards releasing cargo into the extracellular environment or changed activity in sensing cellular amino acid levels (22). There was also an increase in abundance of S-adenosyl homocysteine, SAH, a product of methyl transfer reaction from S-adenosyl methionine, SAM, universal donor of methyl group, to the targets (Supplemental Table 4).

**Figure 5.**
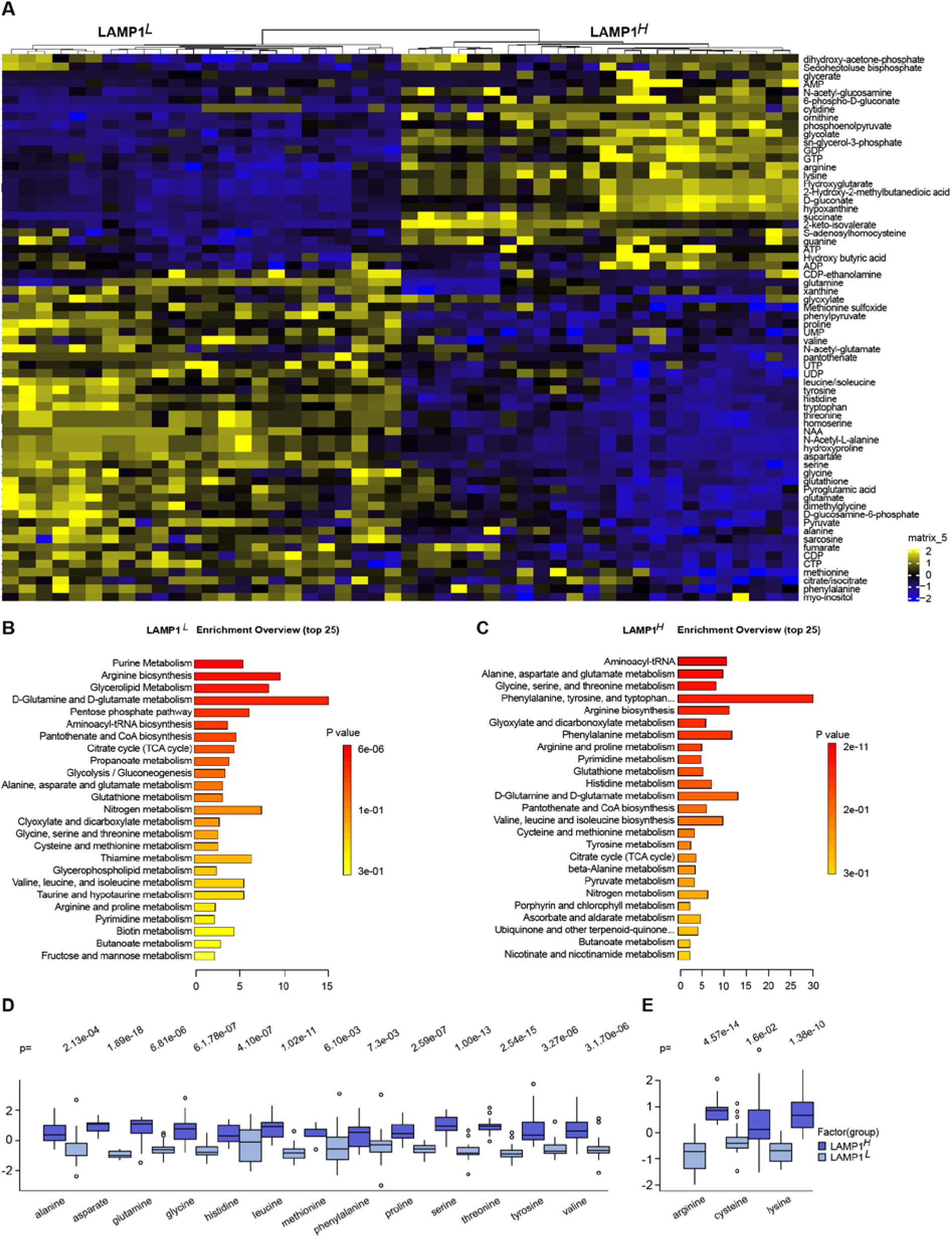
Metabolomic landscape of cells with high LE shows major changes in steady-state levels of amino acids. (A) Heatmap showing differential abundance of steady-state levels of metabolites in LAMP1^*L*^ and LAMP1^*H*^ cells. (B and C) Identification of metabolic pathways from enriched in LAMPL1^*L*^ and LAMP1^*H*^ cells, respectively. (D) Decreased abundance of several amino acids in Lamp1H cells. (E) Increased abundance of three amino acids in LAMP1^*H*^ cells. P values obtained by two-tailed t-test.

### Cells with high LE activity show tumor initiating properties

We have previously shown that knockdowns of LC3C in 786-O VHL(+) and A498-VHL(+) cells induced tumor formation in orthotopic xenografts (15). To test the hypothesis that the tumor forming properties of VHL(+)/LC3CKD cells are related to LE, LAMP^*H*^ and LAMP^*L*^ cells were injected into the flanks of nude mice in a limiting dilution assay. Clearly, cells with high level of LE showed a significantly higher number of tumors at lower number of injected cells, and these tumors were significantly larger than as compared to tumors formed by LAMP1^*L*^ cells (Figure 6A-C). The tumors were histologically malignant, with large areas of necrosis as shown in the H&E staining (Fig. 6D) and strong labeling of the nuclei with proliferation marker, Ki67 (Fig. 6E). The tumor cells maintained high levels of LAMP1 in the plasma membrane as was determined by FACS analysis of cells isolated from individual tumors (Fig. 6F and 6G).

**Figure 6.**
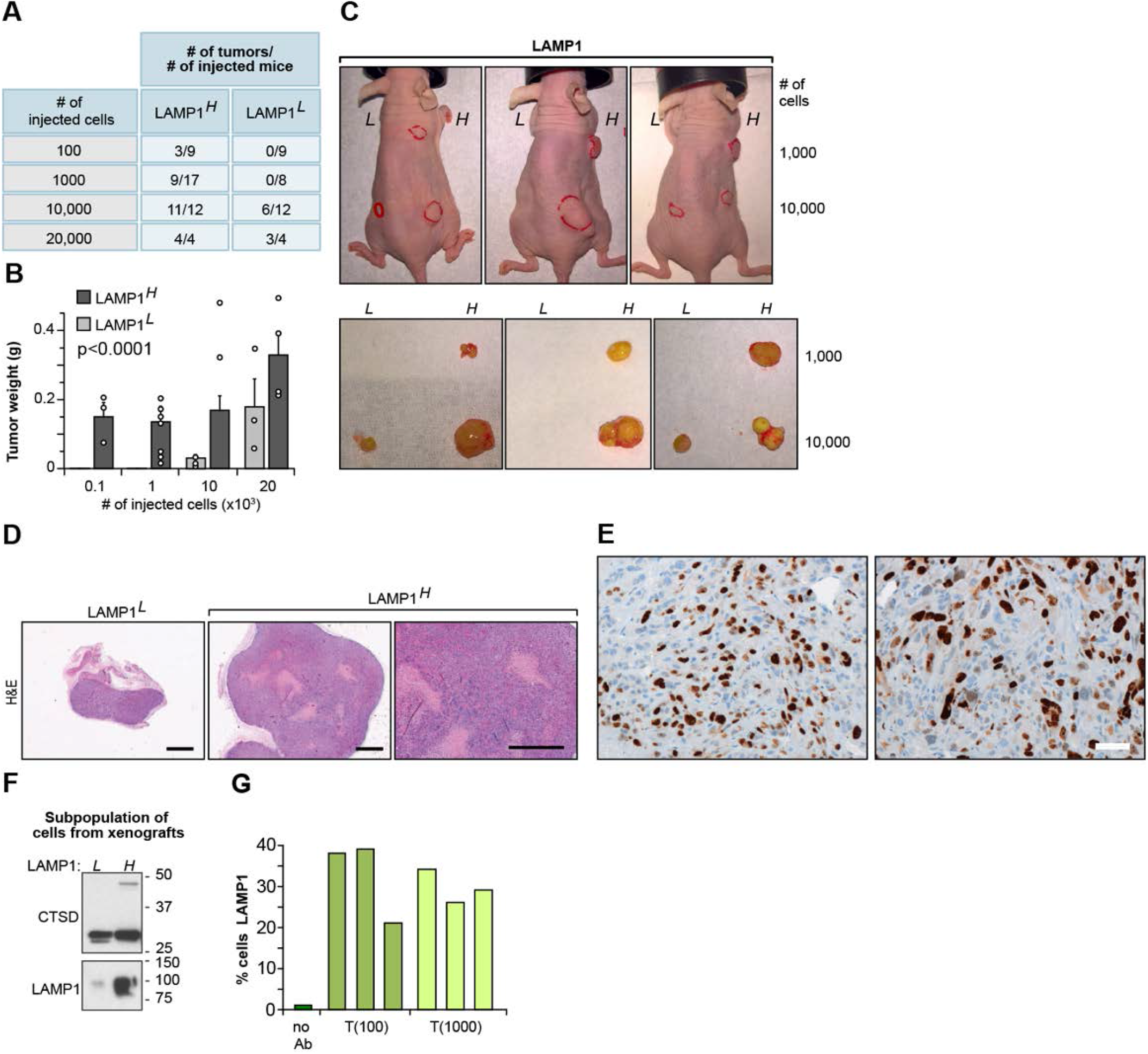
Cells with high, but not low, LE form malignant tumors in xenograft assays. (A) Limited dilution assay. Number of tumors formed by indicated numbers of injected cells. (B) Tumor weight for the indicated numbers of injected cells. (C) representative examples of tumors formed by the indicated numbers of cells. (D) H&E staining of tumors formed by LAMP1^*L*^ and LAMP1^*H*^ cells. Scale bar = 50 µM (E) ICH staining for Ki67 on sections from tumors formed by LAMP1^*H*^ cells. Scale bar = 1 mm (F) Western blot of cells isolated from xenograft tumors maintain high levels of LAMP1 expression. (G) FACS analysis of cells dissociated from three xenograft tumors formed by injection of 100 cells or 1000 cells shows enrichment for cells with LAMP1 present in the plasma membrane.

Analysis of human ccRCC revealed plasma membrane localization of LAMP1 on cancer cells (Fig. 7A), while kidney epithelial cells showed cytoplasmic localization (Fig. 7B). Moreover, analysis of Zn levels revealed significantly lower levels of Zn in ccRCC as compared to the normal kidney tissue (Fig. 7C). These data indicate that lysosomal exocytosis could contribute to lower Zn level in ccRCC and Zn-related epigenetic modifications.

**Figure 7.**
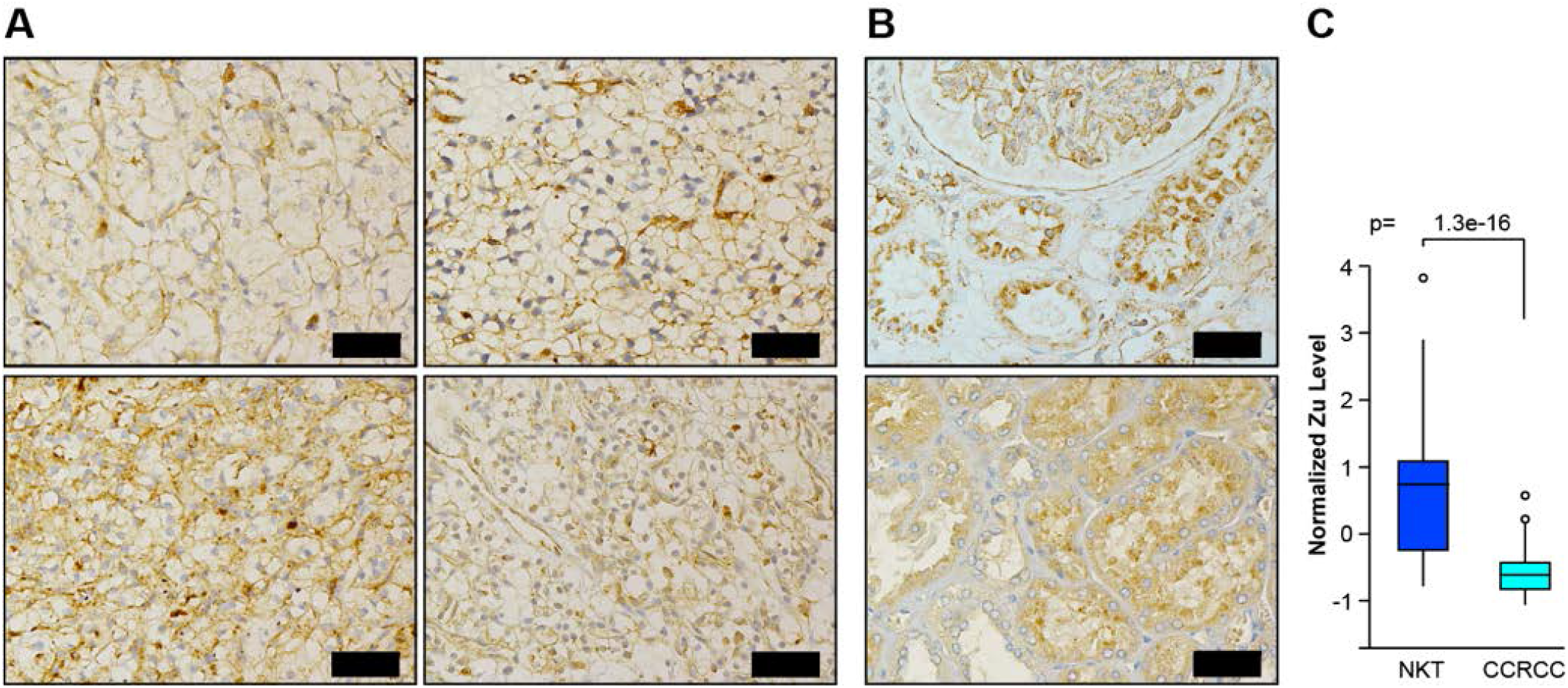
Human ccRCC show plasma membrane localization of LAMP1 in cancer cells and reduced Zn levels as compared to normal kidney tissues. (A) ICH for LAMP1 in sections of 4 ccRCCs. Scale bar = 50 µM. (B) ICH for LAMP1 in sections of two normal kidney tissues. (C) Low level of total Zn in ccRCC as compared to normal kidney tissues. P value obtained by two-tailed t-test.

## Discussion

Lysosomal exocytosis has several important pathophysiological consequences, such as cellular clearance of various cargos and detoxification, membrane repair, and a mechanism supporting cancer cells invasion and metastases. The process requires activity of lysosomal calcium channel, mucopolin 1 (MCOLN1), which releases calcium necessary for the fusion of lysosomal and plasma membrane. Expression of MCOLN1 is under control of TFEB transcription factor, which is also a master regulator of genes involved in lysosome biogenesis and dynamics (23, 24). High expression of TFEB and increased LE are associated with removal of Zn ions, and in turn, TFEB activity is induced by exposure to transition metals (25). Here, we showed that increased presence of lysosomal integral protein, LAMP1, on the plasma membrane is associated with release of cleaved caspase D into the cell culture media. This process was accompanied by translocation of TFEB into the nuclear fraction and induction of gene expression for several lysosomal proteins, including MCOLN1. These data indicate an induction of lysosomal biogenesis in response to augmented LE which impairs intracellular activity of these organelles. Importantly, we found that exacerbated LE caused major change in the intracellular Zn in the intracellular compartments with consequences for epigenetic modification and gene expression. Further on, we determined lower levels of Zn in ccRCC as compared to the normal kidney tissues and presence of LAMP1 on plasma membrane of cancer cells, supporting role of LE in dyshomeostasis of Zn in ccRCC.

Zn is an important trace element with several tumor growth regulating activities (26, 27). It is required for the activity of transcription factors and enzymes containing Zn-finger domains that control transcription and DNA repair, including p53. It has antioxidative properties by regulating expression of metal binding proteins, metallothioneins, and Cu/Zn superoxide dismutase. It regulates cell proliferation and apoptosis. In addition, Zn is likely to affect tumor microenvironment by modulating tumor immunity and inflammation (28). Serum levels of Zn are decreased in several cancers (29). Zn has tumor suppressing activities and its levels are decreased in several solid tumor cancers, including prostate (30, 31), bladder (32), liver (33), bone (34) and head and neck(35). However, in breast cancer, Zn contributes to cancer progression (36). Levels of Zn are regulated by the expression of Zn transporters involved in uptake of Zn (37). Our work indicates that change in lysosomal exocytosis is a powerful mechanism modifying the levels and functions of intracellular Zn.

Importantly, Zn supplementation appears to have positive therapeutic effect in several in vivo animal studies and was suggested in management and chemoprevention of cancer in patients (38). So far information regarding Zn levels in ccRCC has been limited. The low levels of Zn in ccRCC indicate this cancer as a potential candidate for dietary Zn supplementation.

Changes in Zn are known to result in extremely long-lasting epigenetic modifications, including prenatal exposures that epigenetically regulate gene expression in adults (39-41). In particular, Zn deficiency has been shown to decrease histone H3K4 trimethylation and global DNA methylation (41). Here we show that decreased levels of cellular and chromatin fraction of Zn and change in the expression of several Zn regulated proteins correlate with diminished chromatin association of members of PRC2 complex and histone H3K27 trimethylation. The core subunit of PRC2 complex, SUZ12, as well as accessory subunit, MTF2, are zinc-finger proteins. Interestingly, MTF2 was originally described as metal responsive transcription factor, that was able to bind to the metal responsive elements (MRE) in a Zn dependent manner in yeast (21). While ultimately MTF2 was determined to be PCL2 member of PRC2 complexes and to modulate H3K27 methylation, little is understood about the possibility that it could serve as a Zn sensor and translate Zn availability into epigenetic modification regulating gene expression.

The activities of PRC2 are both oncogenic and tumor suppressing (42). High expression of EZH2 is considered to be an unfavorable prognostic factor in ccRCC (43-48), potentially implicating oncogenic function of PRC2 in ccRCC. More in depth analysis of the EZH2 as a negative prognostic factor reveals that it has prognostic value only in males, but not females, and only if stage IV tumors are included in analysis, an indication that its oncogenic activity develops relatively late during cancer progression (The Human Protein Atlas). However, low H3K27 methylation, including H3K27me1, H3K27me2 and H3K27me3, is a negative prognostic factor in ccRCC (49, 50), which would be consistent with our results showing decreased H3K27me3 in cells with high LE that form highly malignant tumors. Clearly, final H3K27 methylation is a readout of combined activities of methyltransferases, demethylases and overall metabolic landscape contributing to the generation of methyl groups. Here it can be proposed that this epigenetic footprint is also regulated by Zn dyshomeostasis.

We found robust metabolic reprogramming related to high LE with potential consequences for epigenetic modifications. The predominant feature, decreased steady-state levels of several amino acids and an increase only in lysine, arginine, cysteine, can be related to altered lysosomal processes, such as degradation of intracellular end extracellular cargos, and can be of consequence in the context of regulation of mTOR activities (22). The amino acid reprogramming can be also causative for the changes in histone methylation. First, there is a decrease in the level of methionine and methionine levels can affect H3K4me3 (51). There is also upregulation of SAH in LAMP1^*H*^. S-adenosyl methionine, SAM, the essential source of methyl groups is synthesized from methionine and ATP by methionine adenosyltransferase. The product of methyltransferase reaction, S-adenosyl homocysteine (SAH), inhibits methyltransferase activity. Thus, small fluctuations in SAM and SAH levels and a decrease of the SAM/SAH ratio can diminish histone methylation at H3K9me2 and H3K27me3 (52). Serine and glycine contribute one carbon units for SAM biosynthesis either by procurement of ATP or methionine recycling (53). Other amino acids, such as threonine, have been shown to be indirectly essential for SAM synthesis and H3K4 methylation in mouse embryonic stem cells (54).

Here we report a new function of LC3C that does not depend on its autophagic activity but rather, potentially on interactions with the microtubules and regulation of lysosomal trafficking (17). LC3C is a microtubule associated protein 1A/1B light chain 3C and in addition to its role in autophagy may promote perinuclear localization of the lysosome, with physiological consequence of supporting fusion of lysosomes and autophagosomes and generation of nutrients. In addition, perinuclear localization of lysosomes leads to decrease in mTOR activity, while peripheral localization increases association of mTOR with lysosomes (55). Movement of lysosomes to the periphery supports tumor growth by remodeling of the extracellular matrix changing the environment for cancer cell adhesion, migration and invasion. Clearly, subpopulation of cells with activated LE shows very aggressive cancer promoting phenotype. Moreover, human ccRCC cancer cells show peripheral positioning of lysosomes as compared to the normal kidney epithelial cells reflecting the role of peripheral distribution of lysosomes in ccRCC progression. The role of LC3C to prevent LE is consistent with its tumor suppressing activity.

## Methods

### Cell culture, treatments, transfection and lentiviral transductions

Human renal cell carcinoma cell lines, 786-O with reconstituted VHL and Caki-1 with endogenous VHL were used. Both cells express high levels of LC3C (14). Stable LC3C knockdowns were performed using GIPZ shRNAs in lentiviral vectors (Dharmacon) (14). For the reconstitutions of LC3C wild type or G126A mutant exogenous LC3C was expressed from the lentiviral construct pLX304 (CCSB-Broad Lenti ORF MAP1LC3C) (14). Lentiviral DNA constructs were VSV-G envelope packaged (Cincinnati Children’s Hospital Medical Center, CCHMC, Viral Vector Core). Cells were grown in DMEM-F12 (Hyclone SH30023) with 10% FBS. All cell analyses were performed 48 h from the original plating. Treatments with Vacuolin (5µM) or chloroquine (20 µM) were performed for 24 h before collection. RNA extractions and QRT-PCR were described before (15). Primer sequences, antibodies and chemicals used are listed in the Supplemental Table 5.

### Flow Cytometry determination of surface LAMP1

Cells were trypsinized, washed with PBS and incubated in DMEM media containing 1% FBS and 1% BSA with Alexa Fluor® 647 Mouse Anti-Human LAMP1 Clone H4A3 (BD Biosciences) for 1h at 4°C with intermittent mixing. Cells were washed in PBS and resuspended in PBS with 1% BSA. Control cells were treated the same way, excluding incubation with the anti-LAMP1 antibody. Xenograft tumors were dissected and minced, following by incubation in F12/DMEM (1:1) with 5% FBS, 20 ug/ml gentamycin, 300 units/ml collagenase and 100 units/ml hyaluronidase at 37^0^C for 2-4 hours. Floating cells were collected and resuspended in prewarmed F12/DMEM with 5% FBS + 2 units/ml DNAse I with antibiotics. Cells were incubated with 30ul LAMP1 ab-Alexa 647, Dapi (1:1000), pacific blue anti-human CD31 (303113, Biolegend) and anti-human CD45 (368539, Biolegend) (2.5 µl/1ml) for 1hr on ice. Filtered single cell populations were sorted and analyzed using a BD FACSAria III. APC conjugated LAMP1 was measured using the 633 excitation laser and a 660/20 nm emission filter. BD FACSDiva software (Version 8.0) was used to run the instrument and to analyze the data. Forward angle scatter, right angle scatter, and fluorescence intensity were recorded from 50,000 cells.

### Immunofluorescence experiments

Cells grown on glass coverslips were fixed with 100% methanol for 5 min at -20°C. Cells were permeabilized with 0.1% saponin. Coverslips were blocked with PBS containing 0.1% saponin and 1% BSA for 30 min and incubated with primary mouse anti-LAMP1 antibody (Abcam, ab25630, 1:3000) or rabbit anti-CTSD antibody (Abcam, ab6313, 1:2000) for 1 hr at 37°C. Coverslips were then washed and incubated with Alexa Fluor labeled secondary antibodies (anti-mouse AF 555 and anti-rabbit AF568, Invitrogen) for 30 min at room temperature. For determination of surface LAMP1 cells were incubated the same anti-LAMP1 antibody (1:50) in PBS with 1% BSA at 4°C for 30 min. Then cells were washed and fixed using 2% PFA and incubated with Alexa Fluor 555 anti-mouse secondary antibodies. Finally, in each case coverslips were washed and mounted using DAPI Fluoromount-G (Southern Biotech, AL).

Confocal images were acquired on a Zeiss LSM710 confocal microscope with a Zeiss Axio.Observer Z1 stand and a Zeiss Plan-Apochromate objective (63x/1.4 Oil DIC) using the Zeiss Zen2010 software. The appropriate lasers and emission filters for the respective fluorophores were used in a multitracking mode. Widefield images were acquired using a Zeiss Axioplan 2 imaging microscope with the appropriate filter cubes and a Zeiss AxioCam MRm B&W camera to record the images using the Zeiss AxioViosion (Rel4.7) software. The objectives had the following specifications: Plan Apochromat 10/0.45; Plan Neofluar 20x/0.5; Plan Neofluar 40x/0.6 (air); Plan Neofluar 40x/1.3 (Oil); Plan Apochromat 63x/a.4 (Oil): alpha-PlanFluar 100x/1.45. All images were saved as 8-bit images.

Quantification of LAMP1 perinuclear vs. peripheral distribution was performed as described (55). Cells were considered to have peripheral vs. perinuclear distribution pattern of lysosomes if more than 50% of LAMP1 staining was in the peripheral vs. perinuclear region, respectively. Approximately 400 cells were counted in three independent experiments.

### Cell fractionation and western blot analysis

Cells were washed, centrifuged and incubated with 5x cell pellet volume of hypotonic buffer A (10 mM HEPES, pH 7.5, 10 mM KCI, 1.5 mM MgCl_2_, 10% glycerol, 1 mM DTT, 0.1% Triton and protease, phosphatase inhibitors) on ice for 5 min. Cells were centrifuged at 1,700g and this step was repeated the second time. The supernatant was collected as cytoplasmic extract. The remaining nuclear pellet was resuspended in 200 ul of high salt buffer B (10 mM HEPES, pH 7.5, 300 mM NaCl, 1.5 mM MgCl_2_, 10% glycerol, 1 mM DTT and protease and phosphatase inhibitors) and extracted with rotation for 1 hr at 4°C. After centrifugation, supernatant was saved as soluble nuclear extract and the remaining chromatin/nucleic acid pellet was washed twice with buffer A to remove salt. Then the pellet was resuspended in 200 ul micrococcal nuclease (MNA)/DNase digestion buffer (50 mM Tris, 50 mM NaCl, 5 mM CaCl_2_. 2.5 mM MgCl_2_) and sonicated 4 times for 20 s. MNA (2 U) and DNAse (37U) were added and digestion was performed for 1 h at 37°C. The reaction w stopped by addition of EGTA to final concentration of 20 mM and centrifuged at 14,000 rpm for 15 min at 4°C to obtain chromatin enriched extract. Cytoplasmic and soluble nuclear extracts were also centrifuged at 14,000 rpm for 15 min at 4°C. Equal protein amounts of each extract were separated on 10% or 4-12% polyacrylamide gels and transferred onto PVDF membrane. Blots were probed with antibodies against: Cathepsin D (abcam ab6313), LC3C (CST 14736), GAPDH (abcam ab8245), TFEB (CST 4240), H3K27me3 (CST 9733), H3K27Ac (CST 4353), Histone H3 (Invitrogen PA5-16183), SUZ12 (CST 3737), EZH2 (CST 5246), MTF2 (Avivia ARP34292 P050), LAMP1 (Santa Cruz sc-5570).

### Xenografts

Experiments on mice were performed in accordance with University of Cincinnati IACUC approved protocols. For subcutaneous injections, indicated number of cells in cell culture medium containing 50% Matrigel were injected into the flanks of athymic nude mice. At the end of 12 weeks, mice were sacrificed and tumors were dissected and weighted and were fixed in 4% paraformaldehyde and processed for H&E staining and immunocytochemistry using anti-Ki67, cell proliferation marker (Mikhaylova et al., 2012). In some cases, tumors were digested to obtain individual cells and surface LAMP1 was determined using Western blot and FACS.

### Zn measurements

Samples were acid digested with nitric acid in a CEM Discover microwave to reduce the carbon load and to mineralize all compounds associated with the elements of interest. Digested samples (1-5 mg) were diluted with ultra-pure water to reduce the acid concentration below 3% and loaded into the ICP-MS-MS (triple quad Agilent 8800x ICP-MS-MS). The instrumental conditions were optimized to remove interferences by using a collision/reaction cell. Integration time was adjusted according to the concentration range for each particular element. Multiple isotopes were monitored when possible to ensure that no interferences were present. The external calibration method was used from 0.01 ng mL^-1^ to 2500 ng mL^-1^ for the elements of interest. A mixture of scandium, yttrium, indium and bismuth was spiked to the samples and calibration as internal standards at 5 ng mL^-1^ to correct for sensitivity drifts. In order to increase the accuracy, internal mass index elemental tags were used in the form of P and S instead of the sample mass. The data analysis was performed with Agilent MassHunter software, with internal standard recoveries and calibration curves. The results are expressed in ng of element per gram of sample. Quality control samples used include NIST SRM 2668-Toxic Elements in Frozen Human Urine standard reference material and the NIST Bovine muscle powder SRM 8414.

### RNA analysis

RNA was extracted using RNAlater ICE (Ambion, AM7030) following by miRNA isolation kit (Ambion, AM1560). The quality of RNA isolated was checked using Bioanalyzer RNA 6000 Nano kit (Agilent, Santa Clara, CA). For RNA seq, polyA RNA was extracted using NEBNext Poly(A) mRNA Magnetic Isolation Module (NEB, Ipswich, MA). RNA-seq libraries were prepared using NEBNext Ultra II Directional RNA Library Prep Kit (NEB). After library QC analysis using Bioanalyzer High Sensitivity DNA kit (Agilent) and library quantification using NEBNext Library Quant kit (NEB), the sequencing was performed under the setting of single read 1×51 bp to generate ∼30 million reads per sample on HiSeq 1000 sequencer (Illumina, San Diego, CA). Data analysis was performed as described before (56).

A list of 8418 Zn-related genes was obtained by querying GeneCards (genecards.org). The overlap of these genes with the LAMP+/-signature was determined (456 genes) and pie charts were generated using R base graphics for both the percentage of Zn-related genes in the LAMP1^*H*^/LAMP1^*L*^ signature and the percentage of Zn-related genes in all protein-coding genes. A chi-square proportions test was performed in R to assess the significance of the proportion of Zn-related genes in the LAMP1^*H*^/LAMP1^*L*^ signature, as compared to the proportion of Zn-related genes as a percentage of protein-coding genes.

For qRT-PCR RNA was purified using Tri Reagent (MRC, TR 118) according to manufacturer’s protocol. cDNA was synthesized using the High Capacity cDNA Reverse Transcription Kit (Applied Biosystems 4368814) with Oligo(dT)20 Primer (Invitrogen 18418020). qPCR was run on a QuantStudio 7 Flex (Applied Biosystems) using 40ng cDNA, Fast SYBR Green Master Mix (Applied Biosystems 4385610), and 400nM primers. Primers are listed in Supplemental Table 5. Fold change was calculated using the ΔΔCt method with GAPDH or 18S as the housekeeping gene.

### Metabolomics

Steady-state of metabolites were measured as described before (56). Cells were homogenized and tissue lysates were mixed 1:1 with ^13^C labeled internal standard mix balanced for the metabolites of interest. The balanced internal standard was generated by combining IROA yeast extract (IROA Technologies) with ^13^C-labelled lysates from several human cell lines grown in the presence of 5.5 mM U-^13^C glucose (Cambridge Isotopes) for three passages. Chromatographic separation was accomplished by hydrophobic interaction liquid chromatography (HILIC) using a Luna NH2 3 µm, 2 mm × 100 mm column, (Phenomenex) on a Vanquish™ Flex Quaternary UHPLC system (Thermo Fisher Scientific). For MS analyses an Orbitrap Fusion™ Lumos™ Tribrid mass spectrometer (Thermo Fisher Scientific) interfaced with an H-ESI electrospray source (Thermo Fisher Scientific) was used. Data were collected for each sample in negative mode using two different mass ranges (70-700 and 220-900 m/z) to enhance sensitivity for larger less abundant compounds, and in positive mode (70-900 m/z). A library of retention times and m/z ratios was compiled using unlabeled and ^13^C-labeled reference compounds and ms/ms fragmentation patterns. These data were referenced to the MZCloud (mzcloud.org) and Mass Bank (massbank.eu) databases. All data were converted to MZXML format using MassMatrix. Peak areas, including isotopically enriched metabolites, were obtained used using Mave for targeted analysis. Differential abundance of metabolites was computed in R using the student’s t-test to determine significance and log2 fold change to determine magnitude and direction of change. Samples were quantile normalized using the R package *preprocessCore* before computing these metrics. FDR corrected p-values were also computed to adjust for multiple-testing bias.

### Human ccRCC specimens and Immunocytochemistry

ccRCCs and normal kidney tissues used for measurements of Zn were de-identified specimens from never smoking Caucasian males described previously (56). For immunocytochemistry sections of fixed and paraffin-embedded tumors were processed in the Pathology Research Core at CCHMC and analyzed by light microscopy. LAMP1 HPA014750 antibody was used (Sigma).

### Statistical analysis

Data are expressed as mean ± SEM for independent experiments ≥ 3. Analysis of differential expression was performed using unpaired two tailed t-test. *P<0.05; **P<0.01,***P<0.001, ****P<0.0001

## Authors contributions

M.F.C-K, J.M, J.L.F. designed the study, supervised the work, and wrote the manuscript; R.V., P.A., M.E.B and B.E. cell culture and flow cytometry experiments. D.S. performed metallomic experiments and analysis; C.W. performed metabolomic experiments. J.R. and J.B. performed bioinformatics analysis of the data. K.VH. provided pathological analysis. X. Z. performed RNA sequencing. JLF supervised metallomic experiments. M.M. supervised bioinformatics analysis of RNA-seq analyses and wrote part of the manuscript. D.R.P contributed to the development of metabolic analyses, writing of the manuscript, and conceptual development of the project.

## Acknowledgments

The work was supported by the following grants: NIH R01GM128216, 2I01BX001110 BLR&D VA Merit, and UCCC Pilot Grant to M.F. C-K. J.M. and M.M were in part supported by UC P30-ES006096 CEG. D.R.P was supported by R01CA168815. C.W. was supported by 5T32CA117846 and ACS PF-17-199. R.V. and P.A. were recipients of American Urological Association Urology Care Foundation Research Scholar Award. We thank Rose Bacon for preparation of the figures.

