## Supplemental Table 5 for "MAP1LC3C regulates lysosomal exocytosis and induces zinc reprogramming in renal cancer cells"

| Supplemental Table 5. Primer Sequences | | |
| --- | --- | --- |
| **Gene** | **Forward Primer** | **Reverse Primer** |
| LAMP1 | TACAATTCTTCCTGACGCGAGACC | TCCGCGTTGCACTTGTAGGAATTG |
| LAMP2 | TGGCAATGATACTTGTCTCTGGC | AGCTGCCTGTGGAGTGAGTTGAT |
| CTSD | TGATTCAGGGCGAGTACA | GGACAGCTTGTAGCCTTTG |
| MCOLN1 | CCACAAGCTGGTCAATGT | TCAGGACGCTGAAGGTATAG |
| NEU1 | TCCAGAGTTCCGAGTGAA | GGTTGCCAGGGATGAATAG |
| TFEB | ATGCCCACCACGCTACC | ATCTGTGAGCTCTCGCTTC |
